# Regulation of Histone Deacetylase 3 by Metal Cations and 10-Hydroxy-2E-DecenoicAcid: Possible Molecular Epigenetic Mechanisms of Queen-Worker Bee Differentiation

**DOI:** 10.1101/415364

**Authors:** Gregory A. Polsinelli, Hongwei D. Yu

**Affiliations:** Department of Biology, Bethany College, Bethany, WV, USA; Department of Biomedical Sciences, Joan C. Edwards School of Medicine, Marshall University, Huntington, WV, USA

**Author notes:** Corresponding author (G.A.P.).

**Keywords:** Histone deacetylase, epigenetics, enzyme kinetics, zinc hydrolase, zinc biochemistry, *Apis mellifera* caste differentiation

## Abstract

Histone deacetylases (HDACs) catalyze the hydrolysis of ε-acetyl-lysine residues of histones. Removal of acetyl groups results in condensation of chromatin structure and repression of gene expression. Human class I, II, and IV HDACs are said to be zinc-dependent in that they require divalent zinc ions to catalyze the deacetylase reaction. HDACs are considered potential targets for the treatment of cancer due to their role in regulating transcription. They are also thought to play important roles in the development of organisms such as honey bees. The fatty acid, 10-hydroxy-2E-decenoic acid (10-HDA), which can account for up to 5% of royal jelly composition has been reported as an HDAC inhibitor. The crystal structure of the HDAC3:SMRT complex possesses two monovalent cations (MVCs) labeled as potassium with one MVC binding site near the active site Zn(II) and the second MVC binding site ≥20 Å from the active site Zn(II). We report here the inhibitory effects of excess Zn(II) on the catalytic activity of histone deacetylase 3 (HDAC3) bound to the deacetylase activating domain of nuclear receptor corepressor 2 (NCOR2). We also report the effects of varying concentrations of potassium ions where [K^+^] up to 10 mM increase HDAC3 activity with a maximum *k*_cat_/*K*_M_ of approximately 80,000 M^-1^s^-1^ while [K^+^] above 10 mM inhibit HDAC3 activity. The inhibition constant (*K*_i_) of 10-HDA was determined to be 5.32 mM. The regulatory effects of zinc, potassium, and 10-HDA concentration on HDAC3 activity suggest a strong correlation between these chemical species and epigenetic control over *Apis mellifera* caste differentiation among other control mechanisms.

## 1. Introduction

The different morphological, reproductive, and behavioral phenotypes observed in the *Apis mellifera* queen and worker bee is interesting considering they are genetically identical. Because they possess the same genome yet display these substantial differences, nutritional control mechanisms are thought to be involved in queen-worker differentiation [1]. These mechanisms are related, in part, to nutritional differences present during development. Queen larvae are fed royal jelly throughout development while worker larvae are fed royal jelly for only the first 1-2 days followed by feeding of worker jelly. Both jellies are a mixture of sugars, amino acids, proteins, fatty acids and minerals. The two jellies have significant quantitative differences [2, 3]. Epigenetic control mechanisms are thought to be modulated by nutritional differences present during queen and worker bee development. DNA methylation appears to play an important role in honey bee caste differentiation and its role appears to be tied to nutrition [4-7]. Another study found that a fatty acid, 10-hydroxy-2E-decenoic acid (10-HDA), present in royal jelly at higher concentrations than in worker jelly reactivated the expression of epigenetically silenced genes in mammalian cells without inhibiting DNA methylation, suggesting 10-HDA is a histone deacetylase (HDAC) inhibitor [8].

HDACs comprise an ancient enzyme family found in plants, animals, fungi, archaebacteria and eubacteria [9]. Histone deacetylases (HDACs) catalyze the removal of acetyl groups from ε-acetyl-lysine residues of histones. Histone acetyltransferases (HATs) acetylate lysine residues of histones thereby activating gene expression. Decreased histone acetylation downregulates affected genes and is associated with cancer development [10]. HDAC inhibitors increase histone acetylation and serve as potential cancer therapeutics [11]. There are at least two FDA-approved drugs, vorinostat (suberoylanilide hydroxamic acid or SAHA) and romidepsin, for treatment of cutaneous T-cell lymphoma with several others in clinical trials [12].

There are four classes of HDACs. Class III HDACs are NAD(+)-dependent and are referred to as sirtuins [13]. This class of HDAC share no sequence similarity with class I and II HDACs and use a different catalytic mechanism [14]. Class II HDACs are subclassified as class IIa (HDAC4, -5, -7, and -9) and class IIb (HDAC6 and -10) and are homologs of yeast HDA1 protein [15, 16]. All members of class IIa can shuttle between the nucleus and cytoplasm. The only class IV deacetylase is HDAC11 [17]. It is a homolog of yeast HOS3. Class I HDACs include HDAC1, -2, -3, and -8 and are homologs of yeast RPD3 [15, 18]. HDACs 1-3 require association with large multisubunit corepressor complexes and are considered inactive by themselves. HDAC8 is fully active in and of itself and is the only extensively kinetically characterized HDAC [19-22].

HDAC3 is unique in that it has a unique domain structure with both nuclear localization and nuclear export sequences [23]. Recombinant HDAC3 cannot be expressed in bacterial cell culture as it is inactive due to improper folding. HDAC3 requires HSC70, TRiC, and most likely HSP90 for proper folding [24, 25]. HDAC3 also requires complex formation with silencing mediator for retinoid or thyroid-hormone receptor (SMRT or NCOR2) or nuclear receptor corepressor 1 (NCOR1) in order to be fully active [26, 27]. The crystal structure of the HDAC3/SMRT shows a channel leading to the active site Zn(II) that is likely obstructed in the absence of SMRT or NCOR1 [28]. Based on the crystal structure of HDAC8, this channel is open offering an explanation for HDAC8 being fully active by itself [29]. It has also been shown that the Zn(II) of HDAC8 can be chelated using EDTA forming apo-HDAC8 and that activity can be recovered by the addition of Zn(II), Co(II), and Fe(II) [19]. It was also shown in the same study that activity was greater for Co(II)-HDAC8 and Fe(II)-HDAC8 and that excess Zn(II) inhibits the Zn(II)-HDAC8, Fe(II)-HDAC8, and Co(II)-HDAC forms.

The catalytic mechanism for HDAC3 and other class I HDACs is based on the crystal structure of histone deacetylase-like protein (HDLP) from *Aquifax aeolicus* [30]. His-142 functions as a general base that deprotonates the metal-activated catalytic water molecule for attack on the substrate amide. A second histidine (His-143) in the active site serves as the acid and protonates the leaving group.

The crystal structures of HDAC3 and -8 show that each bind two monovalent cations (MVCs), likely Na^+^ or K^+^, in addition to the catalytic divalent metal ion [22, 28, 29, 31]. The two MVC binding sites have been designated as site 1 and site 2 with site 1 located approximately 7 Å from the divalent catalytic center and site 2 is ≥ 20 Å from the divalent metal center [28, 29]. The crystal structures of other class I and II human HDACs also bind K^+^ at these same sites in addition to bacterial histone deacetylase-like amidohydrolase [32-34]. A study on the effects of varying concentrations of Na^+^ or K^+^ on catalysis of Co(II)-HDAC8 have been reported [20]. Catalytic activity of Co(II)-HDAC8 was nominally affected by Na^+^ concentration and this MVC was bound to sites 1 and 2 with lower affinity. This study also showed that catalytic activity of Co(II)-HDAC8 is affected by K^+^ concentration to a greater extent with activation of activity up to 1 mM KCl. Higher concentrations of KCl inhibited activity of Co(II)- HDAC8. It was determined that site 1 of HDAC8 is the inhibitory MVC binding site and binding of MVC to site 2 increases activity. KCl concentration was also shown to affect SAHA inhibition of Co(II)-HDAC8 [20].

The present study seeks to determine the effects of excess Zn(II) on catalytic activity of HDAC3 in complex with the deacetylase activating domain (amino acids 395-489) of NCOR2 (or SMRT). We have also examined the effects of varying concentrations of KCl on catalytic activity of HDAC3:NCOR2. The results from these studies confirm and demonstrate that both Zn(II) and K^+^ ion concentrations modulate the activity of HDAC3. We also report the inhibition constant (*K*_i_) of 10-HDA for HDAC3:NCOR2. In the context of honey bee development and caste differentiation, these results suggest a possible link between Zn(II), K^+^, and 10-HDA composition of royal and worker jelly and HDAC activity.

## 2. Materials and Methods

### 2.1. Materials

10-HDA (≥98%) was purchased from Cayman Chemical. All other chemicals were purchased from Sigma Aldrich. Chelex 100 resin was purchased from BIO-RAD. >90% purity (by SDS-PAGE) human HDAC3:NCOR2 in 25 mM HEPES pH 7.5, 300 mM NaCl, 5% glycerol, 0.04% triton X-100, and 0.2 mM TCEP was purchased from Active Motif. All enzyme came from the same lot number. The recombinant complex consists of full length human HDAC3 (accession number NP_003874.2) with a C-terminal FLAG tag and human NCOR2 amino acids 395-489 (accession number NP_006303.4) with an N-terminal 6xHis tag expressed in Sf9 cells. All buffers used in this study were treated with chelex resin prior to use in the enzyme assays.

### 2.2. HDAC3:NCOR2 Activity Assay

The deacetylase activity of the HDAC3:NCOR2 complex was measured using the commercially available Fluor de Lys HDAC3/NCOR1 assay kit from Enzo Life Sciences. Before assaying, HDAC3:NCOR2 was incubated with varying stoichiometric concentrations of KCl. Fluor de Lys-SIRT1 (p53 379-382) substrate (Cat. # BML-KI177) used in the assays comprises an acetylated lysine side chain. Enzyme assays were performed in 96-well plates and reactions were stopped at varying time points using Developer II solution containing 1 μM TSA (an HDAC inhibitor). Fluorescence was measured using a SpectraMax i3x 96-well plate reader with excitation and emission wavelengths of 360nm and 460nm, respectively. Read height was set to 1 mm, 6 flashes per read, and PMT gain set to medium. The concentration of product at each time point was calculated from a standard curve prepared using solutions containing known concentrations of the product (0-40 μM). Except for the zinc inhibition study, all assays were performed in 25 mM Tris pH 8.0 with 500 μM EDTA (free acid) at room temperature.

For Zn(II) inhibition assays, 2 μM HDAC3:NCOR2 was incubated on ice for 1 hour with varying concentrations of Zn(II). The mixture was then diluted to 0.2 μM by the addition of assay buffer and substrate (50 μM) and assayed as above. For assays of HDAC3:NCOR2 dependency on [KCl], varying concentrations of KCl in Tris base, pH 8.0, were incubated with 0.1 μM HDAC3:NCOR2 for 1 hour on ice. In all assays, the final NaCl concentration contributed by the enzyme storage buffer is ≤ 6 mM.

The bell-shaped MVC dependence of HDAC3:NCOR2 on [KCl] was fit by Equation 1 based on Scheme 1 [20]. The concentration of KCl is represented by [MCl], *E*_tot_ is the total enzyme concentration, and the apparent binding affinities for activation and inhibition are represented by *K*_1/2,act_ and *K*_1/2,inhib_. The present study does not analyze concentration effects of NaCl due to its low affinity for MCl binding sites 1 and 2 of HDAC8 accounting for the 100 mM concentration of NaCl needed for full activation of HDAC8 [20]. This study also uses a maximum KCl concentration of 50 mM as it has been shown that the HDAC3:SMRT complex dissociates at salt concentrations exceeding 50 mM [28].

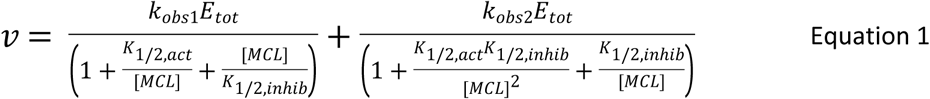

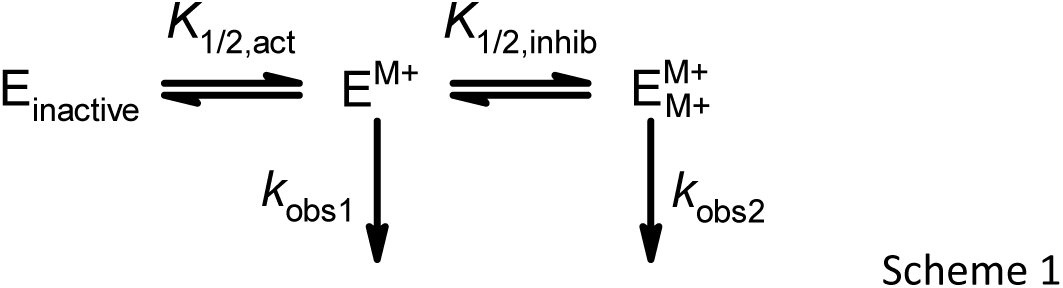

### 2.3. Inhibition by 10-HDA

Inhibition of HDAC3:NCOR2 (5nM final concentration) by 10-HDA (1-10 mM final concentration) was studied by mixing SirT1 substrate (1 μM final concentration) with 10 mM KCl solution and adding this mix to the enzyme on a 96 well plate to initiate the assay. The inhibition constant (*K*_i_) was determined by plotting fractional activity versus [10-HDA] and fitting the data using equation 2. Since the 175mM 10-HDA stock solution was dissolved in 100% DMSO, uninhibited control assays containing the equivalent percentage of DMSO were performed at each inhibitor concentration for fractional activity calculation.

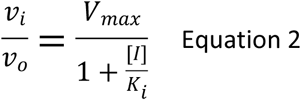

## 3. Results

### 3.1. Zinc Inhibition of HDAC3:NCOR2 Activity

Exhaustive attempts at producing apo-HDAC3:NCOR2 complex via dialysis against the chelators EDTA, dipicolinic acid, and/or 1,10-phenanthroline at various concentrations were unsuccessful. EDTA and dipicolinic acid were successfully used for the preparation of apo-HDAC8 and development of a metal-switching model for the regulation of HDACs [19]. In the present study, dialyzing against EDTA concentrations ≤ 1 mM did not affect HDAC3:NCOR2 activity. At higher concentrations (>1 mM) of EDTA and 1,10-phenanthroline, a decrease in activity was observed (data not shown). However, this activity was unrecoverable upon attempted reconstitution of the treated enzyme using Zn(II) solution suggesting denaturation of HDAC3:NCOR2 and/or dissociation of the complex itself. These observations are supported by refolding studies illustrating the importance of Zn(II) as well as KCl in proper folding of the enzyme [35]. These observations also support the possible role the metal center may play in maintaining a properly folded HDAC3. Based on active site tunnel residues, HDAC3 likely possesses a more hydrophobic environment than HDAC8 preventing access of EDTA to the active site metal.

The addition of 0.1 μM Zn(II) to 50 nM HDAC3:NCOR2 reduced the observed rate 5-fold (Figure 1), indicating a second Zn(II) binding site that is inhibitory of HDAC3:NCOR2 as observed for HDAC8 [19] and other metalloenzymes [36-38]. The lack of linearity observed (Figure 1B) is likely caused by the presence of Zn(II) in the Fluor de Lys SIRT1 substrate. The baseline sample buffer contained 500 μM EDTA which would have chelated contaminating Zn(II) present in substrate and/or elsewhere. Assays with final concentrations of Zn(II) at 0.1 μM, 0.125 μM, and 0.2 μM did not contain EDTA resulting in a greater decrease in activity than expected. An attempt was made to treat the substrate by washing with chelex 100 resin but this resulted in a substantial decrease in fluorescent signal from the treated substrate suggesting the it was bound and removed from solution by chelex resin.

**Figure 1:**
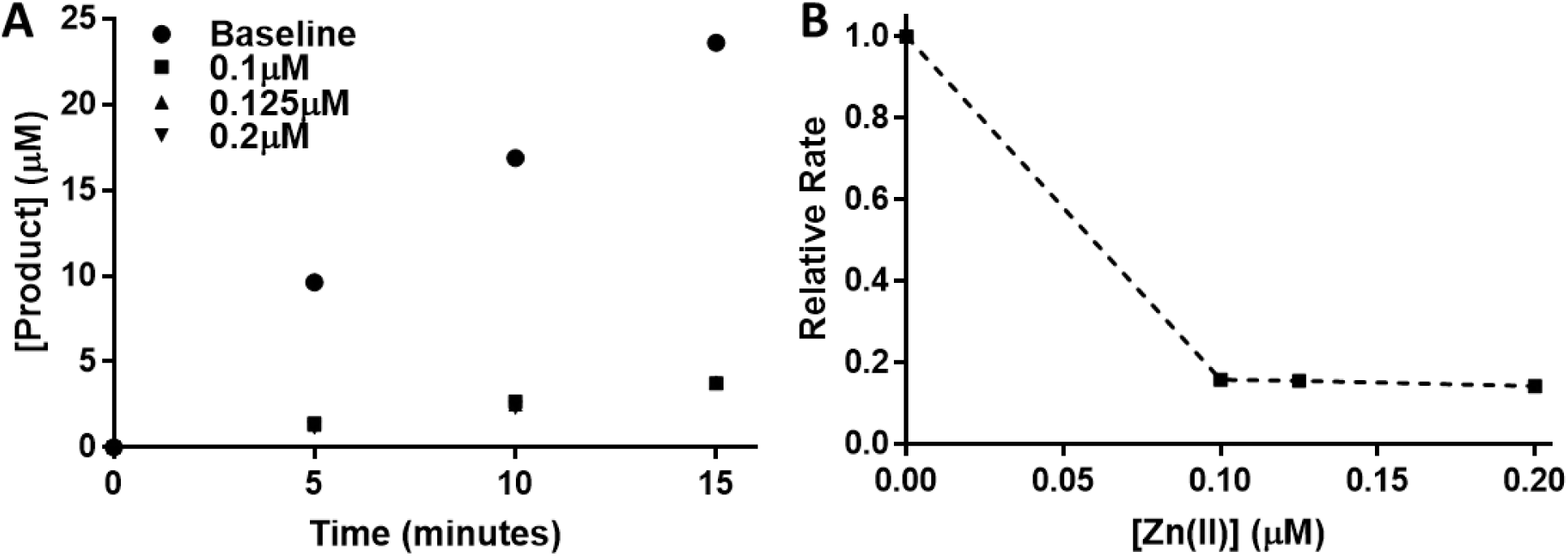
HDAC3:NCOR2 Inhibition by Zn(II). HDAC3:NCOR2 (2 μM final concentration) was incubated with various concentrations of Zn(II) for one hour on ice and then diluted to 50 nM and assayed using the Fluor de Lys assay (panel A). Assay buffer was composed of 25 mM Tris, pH 8, and 10 mM KCl. The baseline sample buffer contained 500 μM EDTA and no added Zn(II). Observed rates at each concentration of Zn(II) were normalized to the baseline rate (panel B).

### 3.2. Potassium Ions Modulate HDAC3:NCOR2 Activity

The roles of the two monovalent cations observed in the HDAC8 crystal structures have been previously studied [20]. However, such a study of HDAC3 has yet to be completed. The activity of HDAC3:NCOR2 and its dependency on KCl concentration was determined by producing Michaelis-Menten plots (Figure 2A) at five different concentrations of KCl to a maximum of 50 mM. The kinetic parameters *K*_M_, *k*_cat_, and *k*_cat_/*K*_M_ were determined by fitting the data to the Michaelis-Menten equation at each concentration of KCl (Table 1). The KCl dependence of HDAC3:NCOR2 is bell-shaped with maximal deacetylase activity at approximately 10 mM KCl (Figure 2B). The data in Figure 2B were fit to equation 1 derived from a two-state sequential binding model (scheme 1) [20]. In this model applied to HDAC3:NCOR2, the enzyme is inactive until the binding of one ion of K^+^ at the activating site. The binding of a second ion of K^+^ results in a decrease in activity. Potassium ion was bound to the higher affinity activation site with a *K*_1/2,act_ of 1.78 mM and to the lower affinity inhibitory site with a *K*_1/2,inhib_ of 46.1 mM. Comparing these values with those determined for HDAC8 [20], HDAC3 shows a two-fold greater affinity for K^+^ binding at the activating MVC site while showing a two-fold decrease in affinity for K^+^ binding at the inhibiting MVC site. From 0.05 mM KCl to 10 mM KCl, K^+^ binding to HDAC3:NCOR2 increased enzymatic activity 140-fold. At its maximum activity, HDAC3:NCOR2 shows an approximate 3.5-fold increase in *k*_cat_/*K*_M_ versus HDAC8. Due to the ionic strength limitations of maintaining the HDAC3:NCOR2 complex, a complete range of KCl concentrations cannot be performed above 50 mM KCl. Therefore, the *k*_cat_/*K*_M_ of the HDAC3:NCOR2 complex with a higher percentage of the two K^+^ ions bound form cannot be experimentally determined. The modulation of HDAC8 activity by NaCl has been determined previously [20]. The maximum *k*_cat_/*K*_M_ was found to occur at 100 mM NaCl. The same limitation prevents a similar study of HDAC3:NCOR2.

**Table 1:**
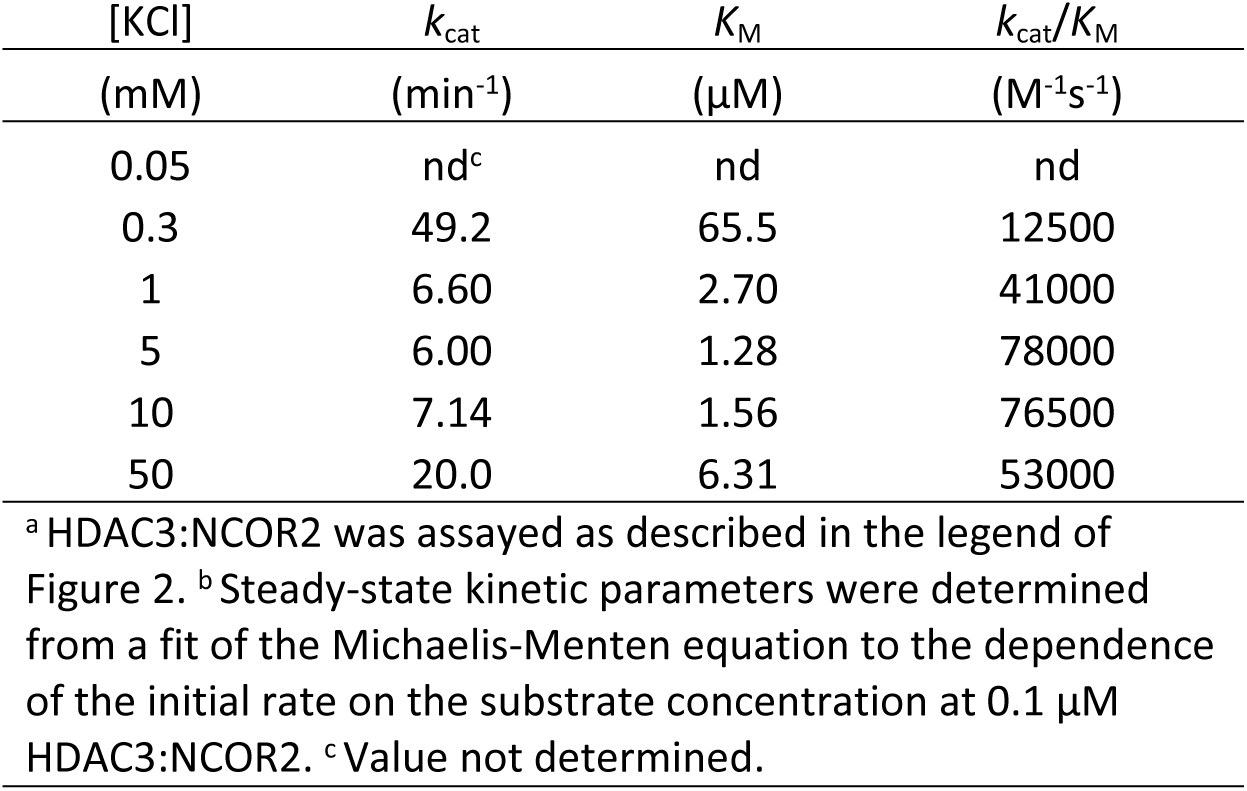
Reactivity of HDAC3:NCOR2 and [KCl] Dependency^a,b^

| [KCl] | $k_{\text{cat}}$ | $K_{\text{M}}$ | $k_{\text{cat}}/K_{\text{M}}$ |
| --- | --- | --- | --- |
| (mM) | (min<sup>-1</sup>) | ($\mu\text{M}$) | (M<sup>-1</sup>s<sup>-1</sup>) |
| 0.05 | nd<sup>c</sup> | nd | nd |
| 0.3 | 49.2 | 65.5 | 12500 |
| 1 | 6.60 | 2.70 | 41000 |
| 5 | 6.00 | 1.28 | 78000 |
| 10 | 7.14 | 1.56 | 76500 |
| 50 | 20.0 | 6.31 | 53000 |

<sup>a</sup> HDAC3:NCOR2 was assayed as described in the legend of Figure 2. <sup>b</sup> Steady-state kinetic parameters were determined from a fit of the Michaelis-Menten equation to the dependence of the initial rate on the substrate concentration at 0.1 $\mu\text{M}$ HDAC3:NCOR2. <sup>c</sup> Value not determined.

**Figure 2:**
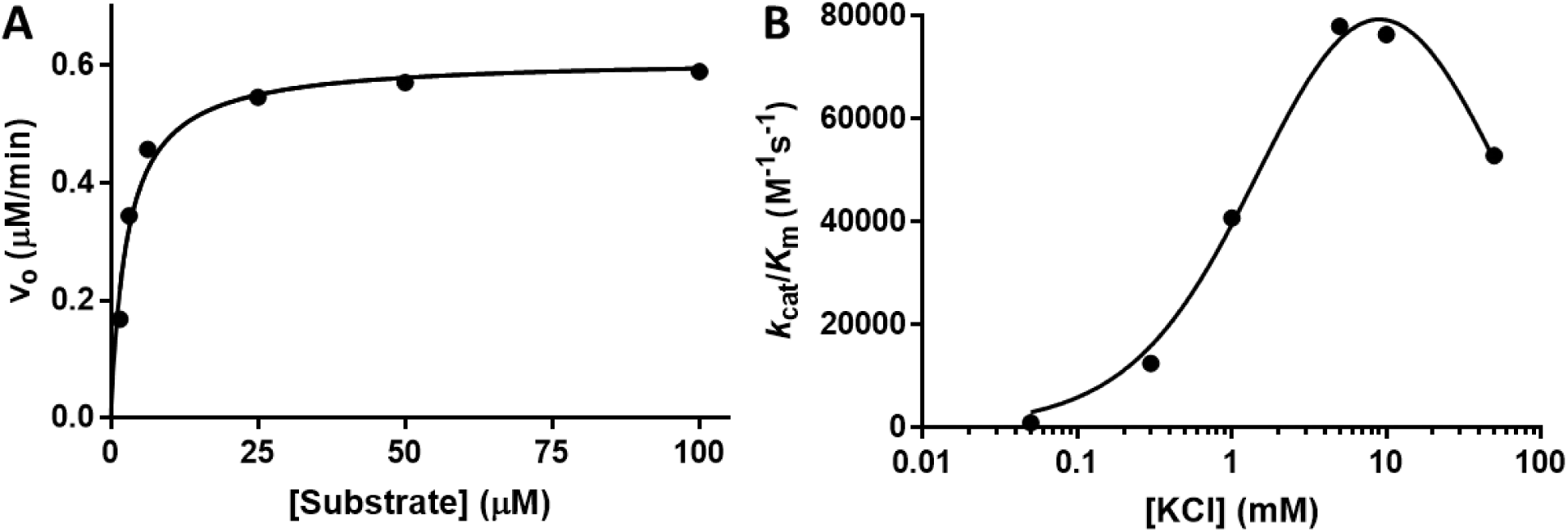
KCl Regulates HDAC3:NCOR2 Catalytic Activity. HDAC3:NCOR2 activity and its dependency on KCl concentration was determined by constructing Michaelis-Menten plots (panel A) at varying concentrations of KCl (0.05 mM-50 mM), Fluor de Lys SIRT1 substrate (1.56 μM-100 μM) and 0.1 μM HDAC3:NCOR2 in 25 mM Tris pH 8.0 with 500 μM EDTA. Initial velocities were determined from time course data based on changes in fluorescence. Catalytic parameters were determined and are summarized in Table 1. Equation 1 was used to fit the bell-shaped dependency of HDAC3:NCOR2 activity with varying [KCl] (panel B) yielding *K*_1/2,act_ and *K*_1/2,inhib_ values.

### 3.3. HDAC3:NCOR2 Inhibition by 10-HDA

The IC50 for 10-HDA and HDAC3 has been reported as 6.5 mM [8]. In the present study, the fractional activity of HDAC3:NCOR2 was determined as a function of 10-HDA concentration and fit using equation 2 (Figure 3). The *K*_i_ from the fit was 5.32 mM and is in agreement with the reported IC50 value. Increasing concentrations of KCl did not significantly affect the *K*_i_ (data not shown). As expected, the results suggest 10-HDA inhibits HDAC3:NCOR2 activity with low affinity *in vitro*.

**Figure 3:**
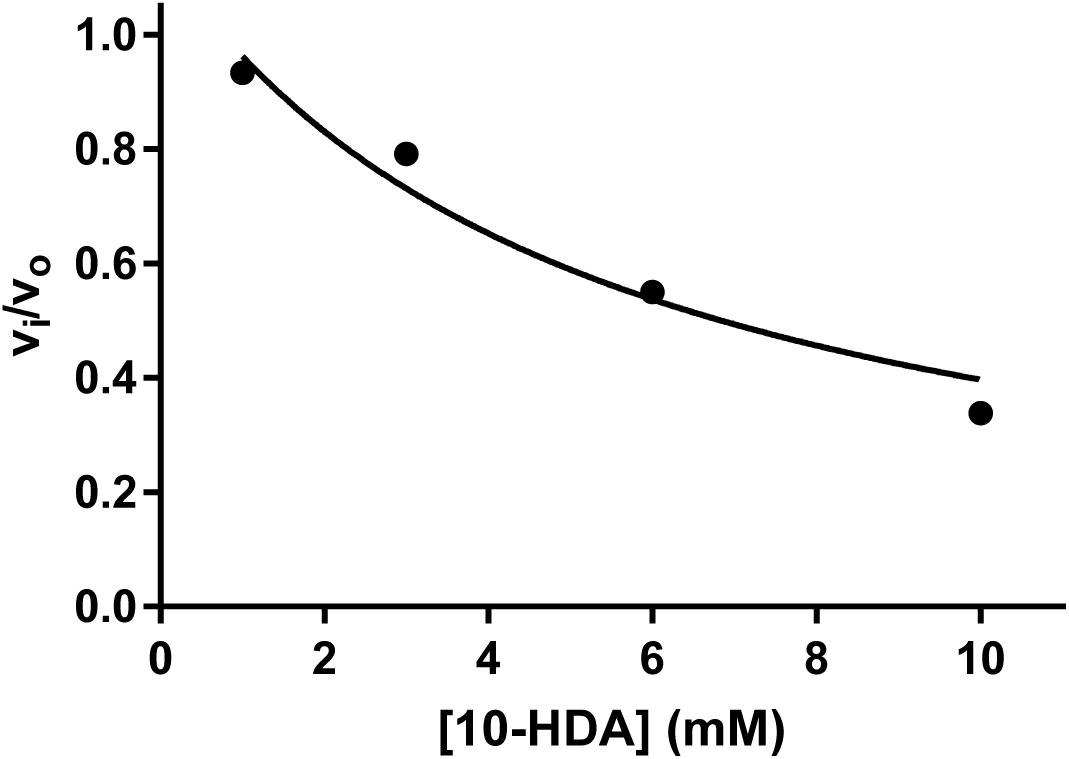
10-HDA Inhibition of HDAC3:NCOR2. A solution containing Fluor de Lys SIRT1 substrate (1 μM final concentration) and 10 mM KCl (final concentration) was added to 5 nM HDAC3:NCOR2 (final concentration) in 25 mM Tris pH 8 with 500 μM EDTA. Initial velocities were determined from fluorescence changes over time, normalized, plotted versus 10-HDA concentration, and fit using equation 2.

## 4. Discussion

### 4.1. Zinc Inhibition of HDAC3:NCOR2

A detailed analysis of metal specificity of HDAC3:NCOR2 was complicated by an inability of metal chelators (EDTA, dipicolinic acid, and 1,10-phenanthroline) to remove the metal center occupying the active site as isolated. HDAC3:NCOR2 activity was decreased 25-50% using higher concentrations of EDTA or 1,10-phenanthroline but this activity was not recovered following reconstitution. Several possible explanations can account for these observations. The metal center is essential to maintain structural integrity of the enzyme and removing it leads to denaturation. Second, the observation that dialysis against 1 mM EDTA and 10 μM dipicolinic acid, conditions that successfully prepared apo-HDAC8 [19], produced no detectable decrease in activity suggest the active site metal is inaccessible to EDTA. The increased active site tunnel hydrophobicity of HDAC3 by comparison to HDAC8 leads the authors to conclude EDTA is unable to access the active site of the enzyme. The expectation was that the less polar metal chelator 1,10-phenanthroline would produce apo-HDAC3:NCOR2. Dialysis against this compound for 24 hours at its maximum solubility in water (15 mM) did not produce the apo form of the enzyme. Attempts at active site metal chelation and removal are ongoing. The identity of the *in vivo* HDAC3 metal is important as the metal occupying the active site has been shown to modulate HDAC8 activity [19]. If Zn(II) of HDAC3 cannot be removed *in vitro*, is it inserted *in vivo* into HDAC3 during folding by another zinc-carrying protein? Can Zn(II) of HDAC3 be replaced via metal-switching as has been demonstrated with HDAC8 and is Zn(II)-HDAC3 the most catalytically active form?

Excess Zn(II) inhibits HDAC3:NCOR2 as has been demonstrated with many other metallohydrolase enzymes [38, 39] including HDAC8 [19]. HDAC8 may not necessarily exist *in vivo* as Zn(II)-HDAC8 leading to the hypothesis of a metal-switching model for HDAC regulation where HDAC8 may exist *in vivo* as Zn(II)-HDAC8, Fe(II)-HDAC8, or even Co(II)-HDAC8 with the Fe(II) and Co(II) forms showing substantially larger *k*_cat_/*K*_M_ [19]. It was also shown in the same study that Zn(II) can inhibit each form of HDAC8. The inhibitory metal binding site on zinc metalloenzymes has been proposed as a potential regulatory mechanism as well [40]. It is interesting to note that the royal jelly fed to queen larvae is significantly higher in zinc content than jelly fed to developing worker bees [3] and the queen is fed royal jelly throughout her life. In addition, the important zinc-binding protein vitellogenin has been positively correlated to high zinc levels, low juvenile hormone, decreased foraging, and longer lifespan in *Apis mellifera* [41-46]. Juvenile hormone has also been positively correlated to stress while vitellogenin protects cells from anti-oxidative damage [45, 46]. We hypothesize that there is a link between dietary zinc levels related to juvenile hormone/vitellogenin titer during development as well as the adult life of *Apis mellifera.* It has been previously shown Fe(II)-HDAC8 can readily be oxidized to an inactive form Fe(III)-HDAC8 [19]. The metal-switching hypothesis for HDACs is appealing based on our knowledge of vitellogenin and juvenile hormone. In the case of high oxidative stress, Fe(II)-HDACs could be readily oxidized to the Fe(III) form resulting in inactive HDAC thereby changing levels of gene expression. In this scenario, vitellogenin and zinc levels are low and therefore foraging activity is high. As oxidative stress increases, levels of juvenile hormone increase resulting in increased foraging behavior and an increase in dietary iron and zinc. The two largest quantities of divalent metals found in worker jelly and royal jelly are zinc and iron with royal jelly possessing significantly greater quantities of zinc than worker jelly [3]. It is also possible that an increased level of Zn(II)-HDAC over Fe(II)-HDAC form would lead to a decrease in oxidative stress and an increase in longevity.

### 4.2. Potassium Modulates HDAC3:NCOR2 Activity

As demonstrated in a previous study with HDAC8 [20], HDAC3:NCOR2 activity is regulated by potassium ions. The crystal structure of HDAC3 in complex with SMRT [28] displayed two bound potassium ions as was observed in the crystal structures of HDAC8 [29, 31]. Like HDAC8, HDAC3:NCOR2 is inactive without potassium and possesses an activation site and an inhibitory site for potassium binding. The activating site of potassium binding has a lower dissociation constant (*K*_1/2,act_ of 1.78 mM) than that of the inhibitory site (*K*_1/2,inhib_ of 46.1 mM). Royal jelly fed to queen larvae has been shown to contain significantly higher concentrations of potassium than that of jelly fed to worker bee larvae [3]. The same study also reported levels of potassium in royal jelly at 3 to 4 times the level of sodium. In the context of the present study, potassium levels in royal and worker jelly likely modulate HDAC activity and levels of gene expression.

### 4.3. 10-HDA Inhibits HDAC3:NCOR2

The IC50 of 10-HDA for several HDACs have been reported in the low mM range [8]. In the present study, the *K*_i_ was determined to be 5.32 mM confirming our expectation of 10-HDA as a weak competitive inhibitor of HDAC3:NCOR2. Royal jelly is composed of 2-5% 10-HDA and is therefore a compelling potential epigenetic regulation factor as previously proposed [8]. Though the IC50 and *K*_i_ are high, the concentration of 10-HDA present in royal jelly is approximately 100mM (at a minimum). A level significantly higher than in worker jelly [3]. Also, the developing queen is fed royal jelly throughout her life providing further support for its role in differentiation and maintenance of health and longevity of the queen.

## Conclusions

HDAC3:NCOR2 is regulated by zinc, potassium, 10-HDA, availability of NCOR1 and NCOR2 as well as inositol phosphates which function in HDAC3:NCOR complex formation [28, 47, 48]. We propose a strong link between queen-worker differentiation, oxidative stress, longevity and dietary levels of zinc, iron, and potassium during the developmental stages and throughout the adult life of *Apis mellifera* based on the modulation of histone deacetylase activity by these chemical species. Based on the complex regulation of HDACs alone, it is unlikely that a single queen determining factor exists but consists of multiple factors that are temporally and behaviorally dependent to include but not limited to the concentrations of zinc, potassium, and iron in some complex regulatory mechanism of modulating vitellogenin and juvenile hormone titer. The possible epigenetic mechanism by which vitellogenin and juvenile hormone expression is controlled requires further exploration. The present study proposes the potential important epigenetic roles of metals present at varying levels in royal and worker jelly leading to plasticity in caste differentiation and behavior.

## Abbreviations

HDAC: Histone deacetylase
SMRT: silencing mediator for retinoid or thyroid-hormone receptor
NCOR: nuclear receptor corepressor
MVC: monovalent cation
DMSO: dimethyl sulfoxide
EDTA: Ethylenediaminetetraacetic acid
10-HDA: 10-hydroxy-2E-decenoic acid
TSA: trichostatin A

